# Topological data analysis quantifies biological nano-structure from single molecule localization microscopy

**DOI:** 10.1101/400275

**Authors:** Jeremy A. Pike, Abdullah O. Khan, Chiara Pallini, Steven G. Thomas, Markus Mund, Jonas Ries, Natalie S. Poulter, Iain B. Styles

## Abstract

The study of complex molecular organisation and nano-structure by localization based microscopy is limited by the available analysis tools. We present a segmentation protocol which, through the application of persistence based clustering, is capable of probing densely packed structures which vary in scale. An increase in segmentation performance over state-of-the-art methods is demonstrated. Moreover we employ persistent homology to move beyond clustering, and quantify the topological structure within data. This provides new information about the preserved shapes formed by molecular architecture. Our methods are flexible and we demonstrate this by applying them to receptor clustering in platelets, nuclear pore components and endocytic proteins. Both 2D and 3D implementations are provided within RSMLM, an R package for pointillist based analysis and batch processing of localization microscopy data.

## Introduction

Single molecule localization microscopy (SMLM) is a super-resolution fluorescence imaging technique capable of localizing individual molecules to approximately 20nm. Since its introduction (1–3), SMLM has matured as a technology and is now routinely used to probe biological nano-structure and processes for a range of biological applications (4, 5). After performing localization, the data from a SMLM experiment is represented by a set of spatial co-ordinates, each corresponding to a single detection, that form a point cloud. This can be analyzed either by rendering an image from these coordinates and using image-based analysis methods, or by analyzing the point cloud directly. Strategies for the latter have focused on the concept of clustering, either by analyzing the spatial statistics of the point cloud to confirm the presence of clustered molecules (6–8), or by grouping individual detections into distinct clusters (6, 9, 10). This latter approach allows per-cluster statistics such as area and detection density to be calculated.

Clustering strategies commonly used for SMLM datasets estimate local detection density and construct clusters from the detections with density above a specified threshold. DBSCAN and Ripley’s K based clustering estimate density using the number of neighboring detections within a specified distance (6, 11), whereas Voronoï diagram based clustering uses the area of the tiles in the associated tessellation (9, 10). The free parameters; a density threshold and sometimes a distance scale, can be set manually or automatically using mean cluster density (9) or Monte-Carlo simulations (10). If assumptions can be made about the distribution and shape of the clusters, a Bayesian engine can be used to set parameters (12). However, biological data is complex, often containing structures of significantly varying density. For such data a single density threshold is not sufficient and a multi-scale approach is required. Clustering algorithms can be repeated, using different parameter values, to segment structures at different densities, for example cells, organelles and protein clusters (9). An alternative approach to density thresholding is to identify clusters based on persistence, or topographic prominence (13). This strategy has shown promise for SMLM datasets in the context of Ripley’s K based clustering (14, 15).

A further limitation of current clustering approaches is that topological information and higher order structure is not considered. Topological data analysis (TDA) provides a robust mathematical framework for probing the topology, or shape, of a point cloud. In this work we employ methods from TDA, specifically persistence based clustering (13) and persistent homology (16–18), to quantify clustering and topological structure within SMLM datasets at a range of scales and densities. We demonstrate their ability to outperform existing methods and reveal new insight into biological nano-structure. Our clustering workflow is used to show a decrease in the area of platelet integrin *α*2*β*1 clusters when the tyrosine kinase Syk is inhibited. Additionally our persistent homology methodology is used to quantify topological structure for endocytic proteins and nuclear pore complex components. The tools are made available to the community as an R package.

## Results

### Persistence based clustering outperforms existing approaches

In common with many existing approaches, the first step in persistence based clustering is the calculation of an underlying density estimate (13). Local maxima within the density estimate are found, and detections assigned to maxima by following the gradient of the density along a specified graph, a collection of points (detections) and lines linking pairs of points. This approach is known as hill climbing and facilitates the separation of clusters in close proximity (19). Next, candidate clusters, or density modes, are merged based on the strength, or persistence, of each local maximum. For each mode the persistence is defined as the difference between the local maximum; the birth density, and and the corresponding local minimum; the death density. This clustering scheme is named the Topological Mode Analysis Tool (ToMATo) and is analogous to local thresholding of the density estimate (13). Therefore ToMATo is capable of segmenting clusters at varying densities simultaneously.

To evaluate ToMATo on SMLM data we generated realistic simulated dSTORM datasets of Gaussian clusters. Low, high and mixed (a mixture of high and low) density clusters were generated, either in close proximity or well separated (**Supplementary Fig. S1**). Local detection density was estimated by counting the number of other detections within a fixed radius, and candidate clusters were constructed from a graph linking all detections within the same search radius (**Fig. 1a,b**). With our implementation of ToMATo there are two free parameters; the search distance and a threshold on cluster persistence. To enable the selection of a suitable persistence threshold a scatter plot of the death and birth densities for each candidate cluster can be plotted. This is known as a ToMATo (persistence) diagram (**Fig. 1c**). Candidate clusters below the persistence threshold are weak and are merged to neighboring clusters (**Fig. 1d**). Detections which cannot be linked to a cluster above the persistence threshold are considered to be noise.

**Fig. 1.**
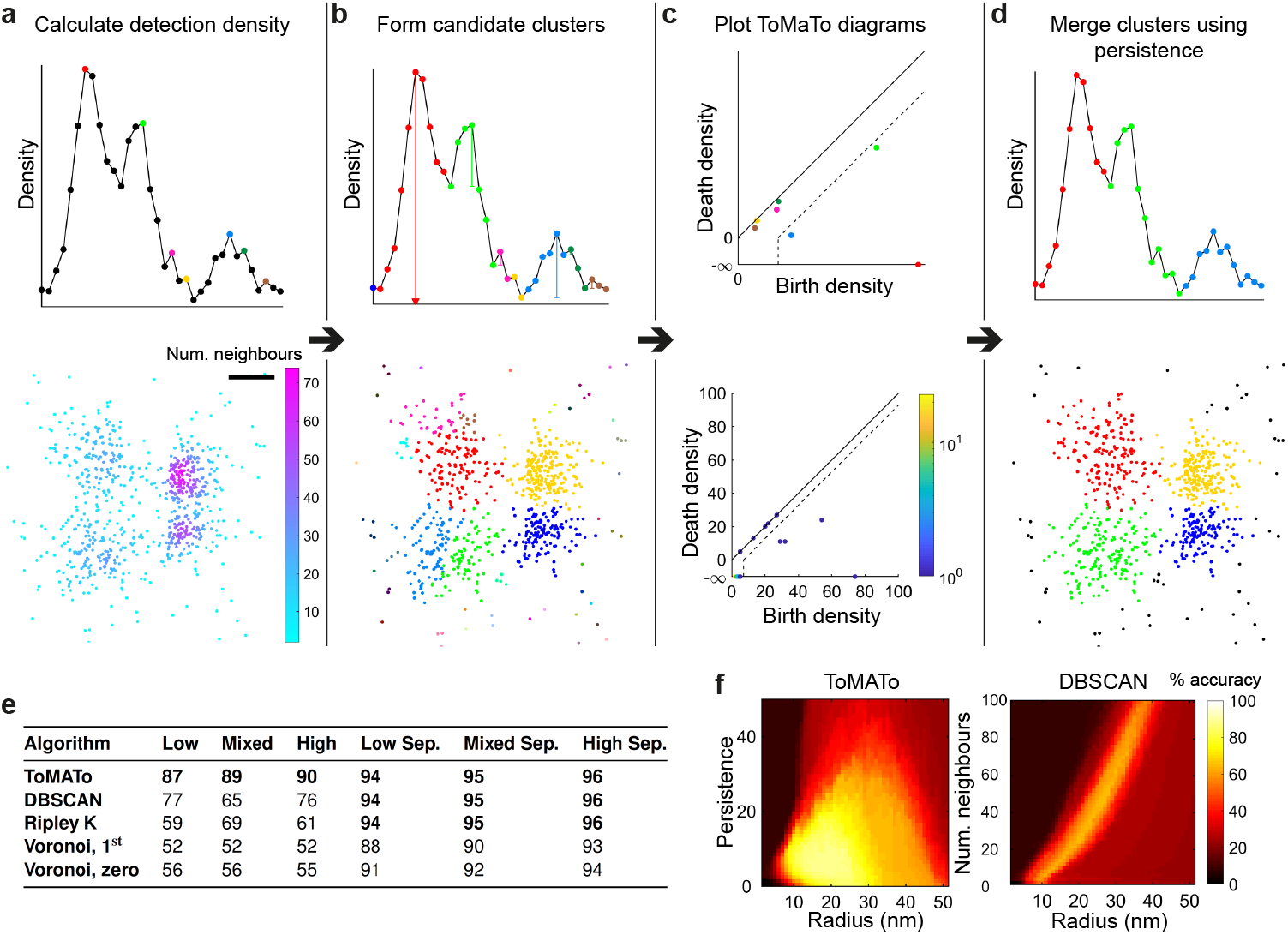
Persistence based clustering for SMLM data. (**a**) The first step is the calculation of detection density. Top row shows an illustrative 1D example with three prominent clusters. Bottom row shows a 2D dSTORM simulation of four Gaussian clusters in close proximity with unequal variance (mixed density). Here density is estimated by counting the number of neighboring detections within a fixed search radius, set to the optimal value of 19nm. Scale-bar 50nm. (**b**) Detections are assigned to local maxima in the density estimate by following the gradient of a graph formed by linking detections within the same search radius. These density modes form candidate clusters. (**c**) ToMATo diagrams showing the birth and death density for each candidate cluster. The vertical distance from the diagonal represents the persistence of the candidate and a threshold is chosen below which clusters are merged (dotted line). For the dSTORM simulation this is set to the optimal value of 6 detections. The highest peak in each connected component of the graph never dies and resides at death density *−∞*. (**d**) Final ToMATo clustering results after cluster merging based on persistence. Noise detections, shown in black, are formed when detections cannot be merged to a persistent cluster. (**e**) Performance of clustering algorithms was quantified as the percentage of correctly assigned detections. Six different scenarios were simulated: low, mixed and high density clusters either in close proximity, or well separated (Sep.). For each scenario twenty simulations were analyzed and the maximal performance (averaged across simulations) for all parameter sets is shown. (**f**) Performance of ToMATo and DBSCAN across all tested parameters for the mixed density dataset.

We compared persistence based clustering to existing routinely used approaches, specifically DBSCAN (11), Ripley’s K based clustering (6, 12), and Voronoï tessellation (9, 10). A range of free parameters were used for each simulation scenario and algorithm. Performance was quantified as the percentage of correctly assigned detections and averaged over repeated simulations (**Supplementary Fig. S1**). ToMATo significantly outperforms these existing approaches in challenging scenarios when clusters are close together (**Fig. 1e**). For the easier simulations where clusters are well separated it performs equally well. Moreover ToMATo is less sensitive to small changes in the choice of free parameters (**Fig. 1f** and **Supplementary Fig. S2**). Together these results show that persistence based clustering is the highest performing and most stable of the tested algorithms for SMLM cluster analysis.

### Syk inhibition reduces the area of integrin *α2β1* clusters in platelets

To demonstrate the use of persistence based clustering on SMLM datasets we segment nano-structures of integrin *α*2*β*1 in platelets seeded on collagen fibres (**Fig. 2a,b and Supplementary Fig. S3**). Integrin *α*2*β*1, a platelet collagen receptor, accumulates at collagen fibres in spread platelets as shown in **Supplementary Fig. S3**(20). Within a single platelet there are areas with sparse or tightly packed *α*2*β*1 clusters due to differences in the underlying collagen distribution. This therefore represents a difficult multi-density segmentation problem. Stable platelet adhesion to collagen via *α*2*β*1 under flow conditions is dependent on the presence of the cytoskeletal adapter protein talin, which links integrins to the actin cytoskeleton (21, 22). We disrupted the cortical actin organisation using the tyrosine kinase Syk inhibitor PRT060318 (23, 24), which we hypothesised would interfere with *α*2*β*1 clustering on collagen fibres. Syk inhibition results in a significant reduction in mean cluster area, however, no significant difference in cluster density was observed (**Fig. 2c**).

**Fig. 2.**
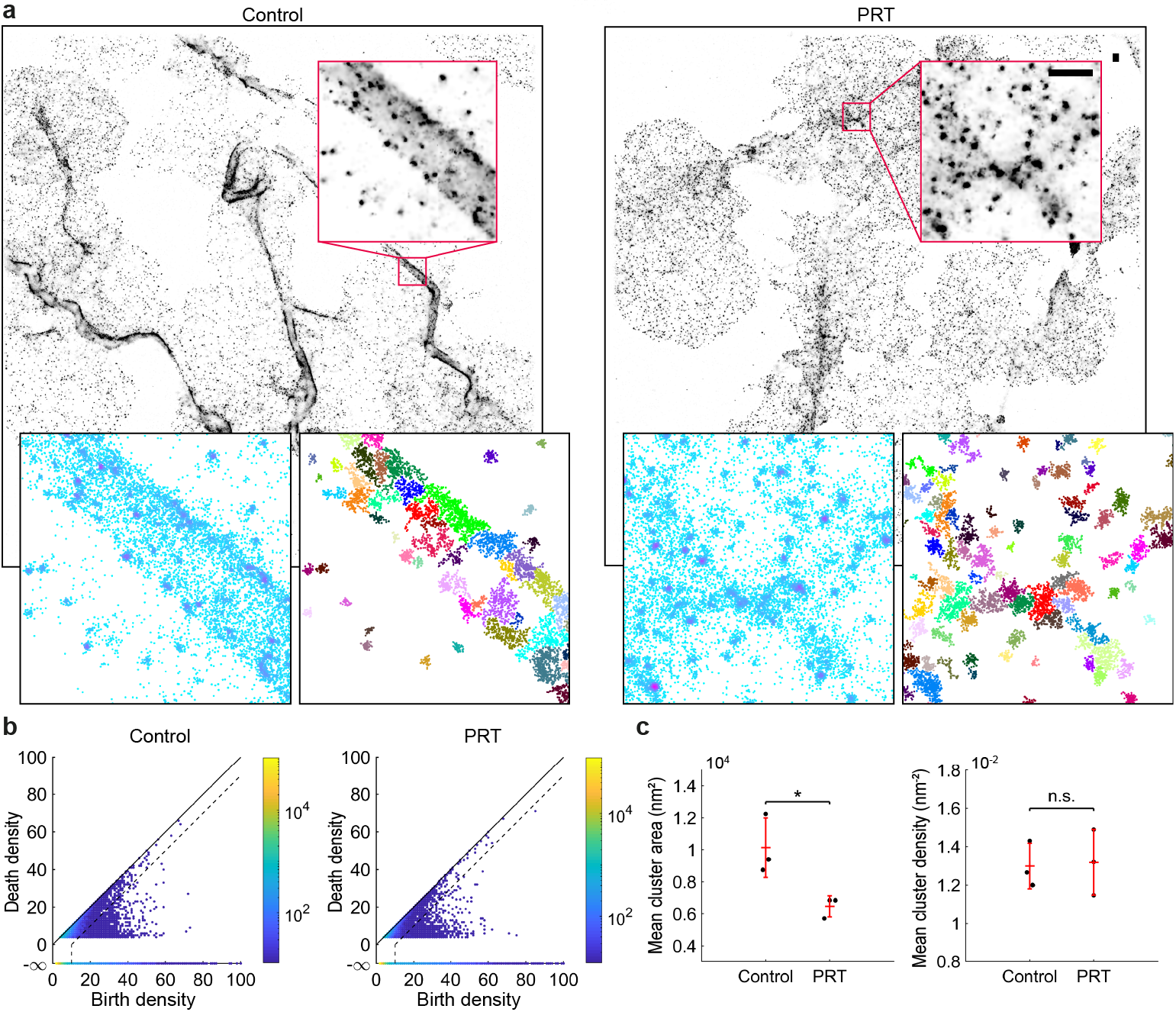
Syk inhibition reduces the mean area of integrin *α***2***β***1** clusters. (**a**) Platelets were seeded onto collagen fibers and treated with either the Syk inhibitor PRT060318, or a DMSO control. The sample was immunollabeled for integrin *α*2*β*1, secondary labelled with AlexaFluor647, and imaged using dSTORM. Persistence based clustering (ToMATo) was used to segment integrin *α*2*β*1 nano-structures. Representative dSTORM image reconstructions, density estimates and clustering results (noise not shown). The search radius for the calculation of the density estimate and linking graph was set to 20nm. Scale-bar 500nm. (**b**) ToMATo diagrams showing the birth and death density for each candidate cluster. Dotted line shows the chosen persistence threshold for merging of clusters (10 detections). (**c**) Mean cluster area and cluster density. N = 3, four fields of view per replicate. The entire field of view was analysed and mean cluster statistics were computed for all clusters in a replicate. Comparisons by two-sample t-test (^*^*P* < 0.05), error bars are mean *±* s.d.

### Persistent homology quantifies topological nano-structure

SMLM data is information rich and contains more structural insight than is available through cluster analysis alone. Here we use persistent homology to extract complementary topological information from the data. The concept of a graph can be extended to a higher dimensional structure, known as a simplicial complex. A simplicial complex is constructed from points, lines, triangles, tetrahedrons and equivalent higher order structures, collectively known as simplices. For SMLM data we build the simplicial complex on top of the point cloud formed by the detection list. There are a variety of methods for building complexes but this work focuses on the Rips complex, an abstract simplicial complex chosen for it’s efficient computation and storage. If each point within a candidate simplex (two points for a line and three for a triangle) is within a search distance of every other point, then the simplex is included in the complex. From the Rips complex topological features of the underlying point cloud can be computed as a function of search distance, or scale. First order features correspond to the number of connected components in the complex, second order to the number of *holes* or *loops*. When working with 3D datasets, the third order features are enclosed *voids*.

Computing the topological structure within the data at a single scale is not very informative; any given feature could be unstable due to small variations in scale and it is not possible to capture multi-scale structure. To overcome this the Rips complex is computed for a range of scales, a process known as a filtration (**Supplementary Video 1** and **Fig. 3a,b**). To summarise the information present in a filtration, the birth scale (emergence) and death scale (closure) of each topological feature is plotted in a persistence diagram (**Fig. 3c**). The more robust a feature is to changes in scale, the longer it persists, and the greater the distance from the diagonal of the diagram. Fragile features, typically noise, will be located close to the diagonal. Therefore thresholding features by persistence selects only the most robust, a procedure known as persistent homology (16–18).

**Fig. 3.**
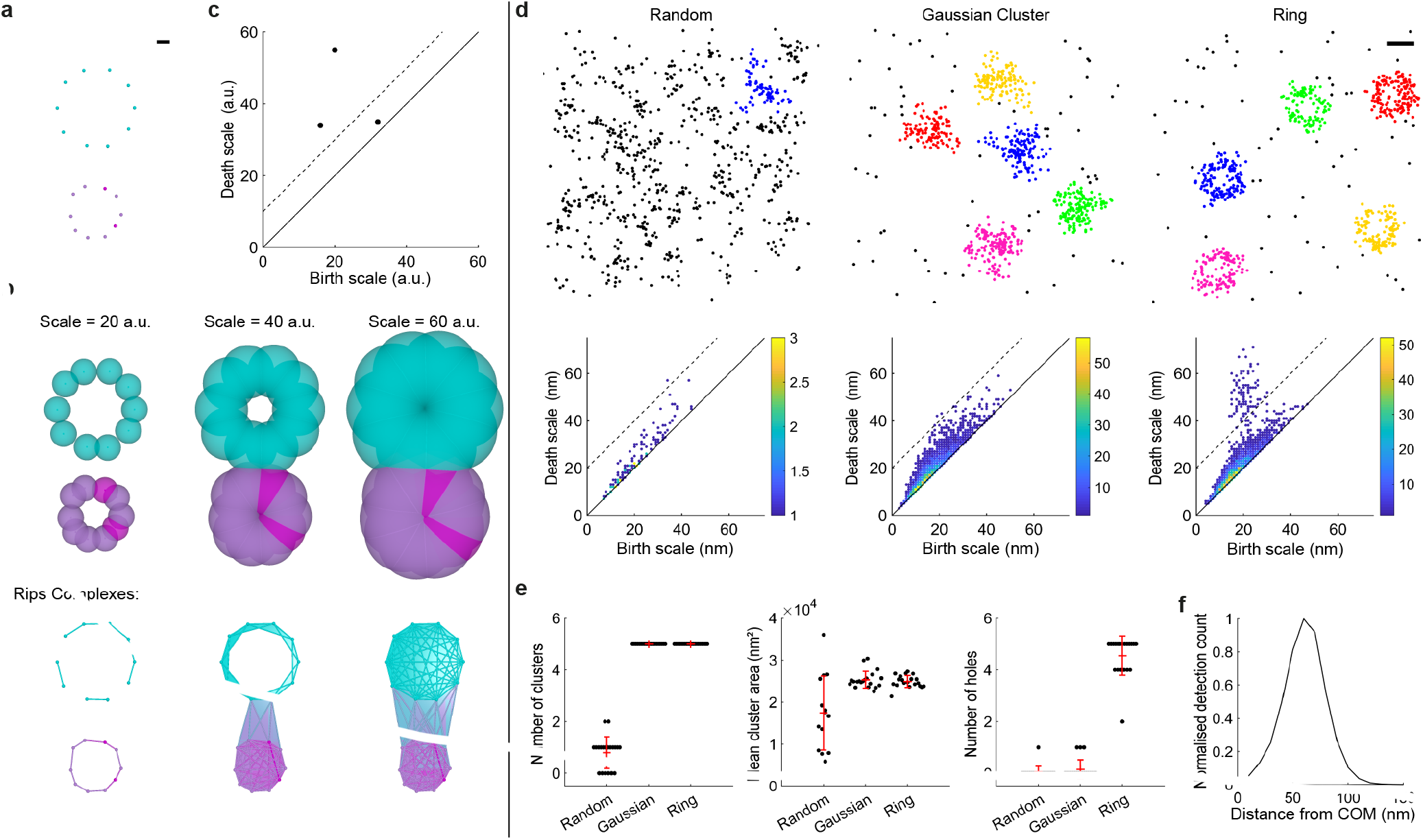
Persistent homology for topological analysis of clusters in SMLM datasets. (**a**) Illustrative example where detections are spaced evenly on the circumference of two circles. Scale-bar 10 arbitrary units (a.u.). (**b**) Building a filtration. Balls of varying diameter were placed at each detection (top) and the Rips complexes (bottom) were determined by the overlap of these balls. The filtration was evaluated for all integer values between 1 and 60a.u.. Without the filtration, it would be difficult to choose a scale which fully encapsulates the clustering and topology of the data. Simplex colour is set by the detection density estimate and is only for display purposes. (**c**) The persistence diagram summarises structure within the filtration. The birth and death scales for each hole are shown. The persistence threshold is shown as a dotted line and there are two significant holes above this threshold. (**d**) Simulations for randomly distributed molecules, Gaussian clusters and rings with 60nm radius were segmented using ToMATo. For each cluster a filtration was constructed and the corresponding persistence diagrams are shown. All holes have been grouped into a single persistence diagram per scenario. A persistence threshold of 20nm was applied to all scenarios (dotted line). Scale-bar 100nm. (**e**) Mean number of clusters, cluster area and number of holes. Error bars are mean *±* s.d.. (**f**) Averaged radial distribution for all clusters with a single hole from the ring simulation. As expected the peak of the profile lies at 60nm.

To test our persistent homology based workflow for SMLM data, realistic synthetic datasets were generated. Molecules were distributed according to one of three scenarios; (i) complete spatial randomness (CSR), (ii) Gaussian clusters, or (iii) circular rings with molecules evenly distributed on the circumference. Before performing persistent homology it is advantageous to perform a cluster analysis to segment the data into nano-structures (**Supplementary Fig. S4**). This reduces the effects of noise, and also the computational cost. We used ToMATo to segment the clusters and subsequently grouped topological features across all clusters into a single persistent diagram per scenario (**Fig. 3d** and **Supplementary Video 2**). A single persistence threshold was applied to all scenarios and the number of significant holes, for each cluster, was counted (**Fig. 3e**). ToMATo clustering was able to accurately segment clusters and persistent homology showed that the number of holes was significantly higher for the ring simulation, and in most cases equal to the number of clusters. As expected no significant difference in cluster area was found between Gaussian and ring clusters. Therefore standard cluster statistics cannot distinguish between Gaussian and ring clusters, whereas persistent homology clearly reveals the key topological difference. Several clusters were found in the CSR scenario due to artifacts such as multiple blinking events. However the number of holes for these clusters was not significantly greater than zero.

The topology of clusters, as defined by our workflow, can be used to filter clusters for downstream analysis. For example we can select all clusters with a single hole and find the average radial distribution as shown for the ring simulations in **Fig. 3f**. Our workflow can also be applied to 3D SMLM datasets as demonstrated for equivalent simulations in **Supplementary Fig. S5** and **Video 3**.

### Topological analysis of nuclear pore and endocytic proteins

To evaluate our novel methods on real data we focused on structures for which the topology has already been well characterised through image based particle averaging. This is appropriate as it facilitates a robust validation of the proposed workflow. Specifically we choose Nup107, a component of the nuclear and cytoplasmic rings of nuclear pore complexes (25–27), and three different proteins of the yeast endocytic machinery; Las17, Ede1 and Sla1 (4).

Persistence based clustering was used as a pre-processing step to segment either the nuclear pore complex, or endocytic sites (**Supplementary Fig. S6, S7, S8**). For Nup107 both 2D and 3D dSTORM datasets were analyzed. For the 2D dataset the resulting persistent diagram shows a large number of holes above the threshold (**Fig. 4a**). As expected most (65%) of the clusters have a single hole topology. 32% of clusters have no significant holes (**Fig. 4b**). These clusters could be a result of many scenarios including clustered molecules not in a pore complex, imaging artifacts, noise detections in the center of features and variation in pore alignment. Filtering of clusters with a single hole results in an average radial profile with a peak at 60nm, reproducing the results of imaged based particle averaging (**Fig. 4c**) (27). When imaged in 3D both the cytoplasmic and nuclear rings of the complex must be considered. If the cluster segmentation is able to split these two rings each cluster will have a single hole. If a cluster contains both rings the resulting structure will have two holes and no enclosed voids. When our method is applied to 3D data we observe large numbers of structures with either single (36%), or two (16%) holes (**Supplementary Fig. S8**). Very few structures had enclosed voids (2%). Again there is a large number of structures with no topological features (39%).

**Fig. 4.**
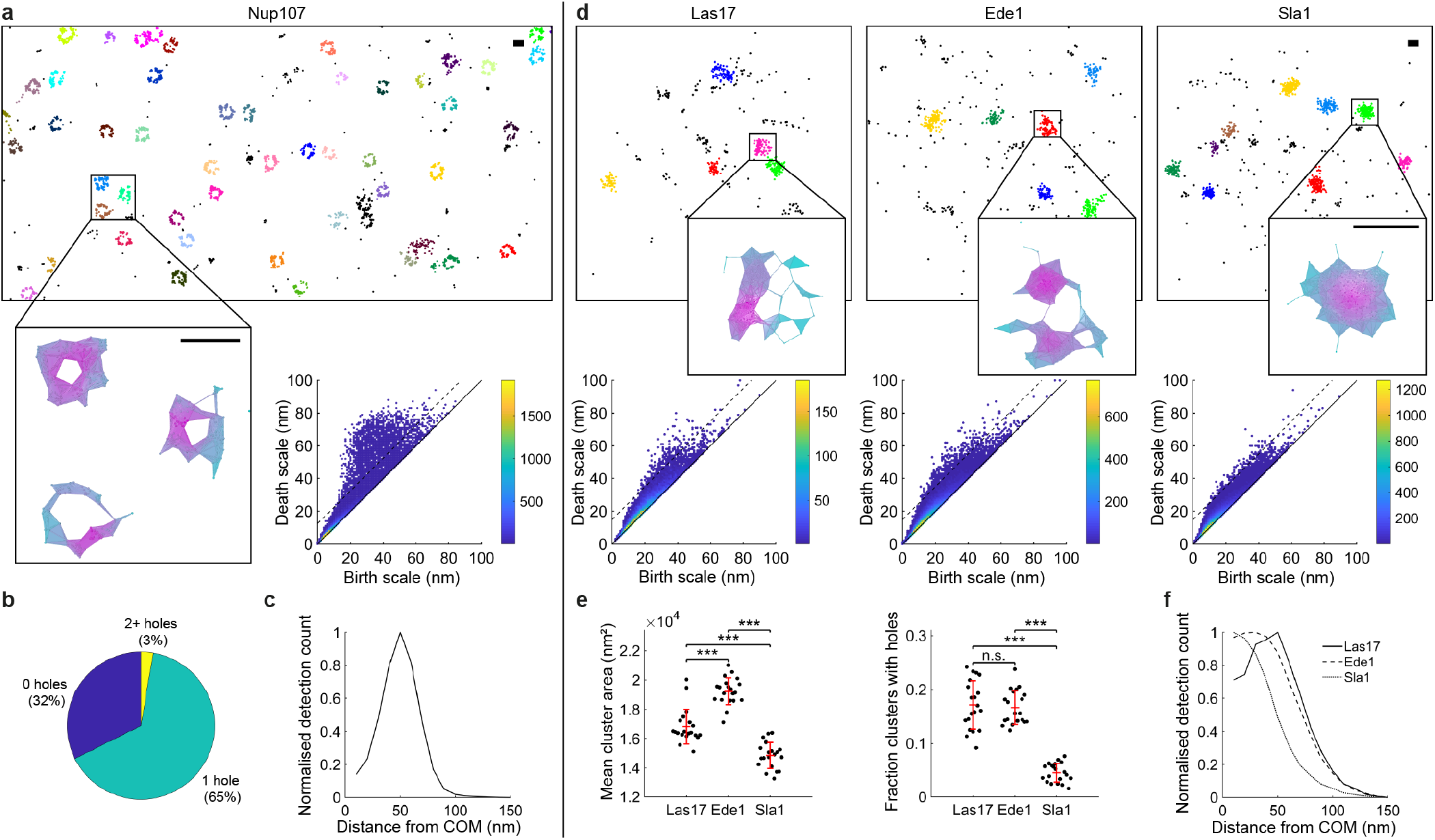
Persistent homology quantifies the topological configuration of biological nano-structures. (**a**) Topological analysis of Nup107-Snap-AlexaFlour647 imaged using dSTORM. Cropped field of view showing clustering result and example Rips complex evaluated at 40nm. Scale-bar 100nm. A threshold of 15nm was chosen for the persistence digram (dotted line). (**b**) Percentage of Nup107 clusters with specified topological configuration. (**c**) Averaged radial distribution for all clusters with a single significant hole. Peak at 50nm. (**d**) Topological analysis of the endocytic proteins Las17, Ede1 and Sla1 in yeast. Cropped fields of view showing clustering results and example Rips complexes evaluated at 30nm. Persistence threshold was set to 15nm (dotted lines). (**e**) Mean cluster area and the percentage of clusters with significant holes. Twenty fields of view were analysed. Statistics performed by one-way ANOVA and subsequent tests for multiple comparisons using the Bonferroni method (^***^*P* < 0.001). Error bars are mean *±* s.d.. (**f**) Averaged radial distributions for all clusters with a single significant hole. Peak for Las17 profile at 50nm.

In a recent study (4) the actin nucleation promoting factor Las17, has been shown to have a clear ring profile when many endocytic sites are averaged, whereas the coat protein Sla1 does not. It was also shown that Ede1 is recruited to sites in the early staged of endocytosis, and clusters are not uniform in size and shape, hence a ring like topology is not clear from image based particle averaging. We take data from this study, where endocytic proteins in yeast were endogenously labeled with a photoconvertible fluorescent protein and imaged using a homebuilt SMLM system, and apply our novel topological analysis workflow. Figure **Fig. 4d** shows example cluster results and persistence diagrams for all three proteins. From visual inspection of the diagrams it is clear that both Las17 and Ede1 clusters have large numbers of holes, whereas Sla1 clusters do not. Quantification of the percentage of clusters with holes reveals a significant difference between Las17 and Sla1, and Ede1 and Sla1, but not Las17 and Ede1 (**Fig. 4e**). The averaged radial profile of single hole clusters for Las17 has a clear ring profile with maximum at 50nm, confirming the results of imaged based analysis (**Fig. 4f**). Together these results demonstrate that our method is able to accurately and efficiently quantify nano-scale topology in SMLM datasets.

### RSMLM: R package for analysis of SMLM datasets

To complement this study we have released RSMLM, an R package for the pointillist based analysis of SMLM data. This package includes the methods described in this paper for persistent clustering (ToMATo), alongside DBSCAN, Voronoï tessellation and Ripley’s K based clustering. There is also the capacity to simulate dSTORM data. This library will provide an adaptable framework for analysing and batch processing both 2D, and 3D, SMLM datasets. Binder ready Jupyter notebook tutorials are provided to facilitate easy use of the package. The functionality of the library can also be included within KNIME workflows using simple R-snippets (28). This enables users without any scripting knowledge to access the core functionality.

## Discussion

As single molecule methods for high throughput data acquisition (29) and sample labelling (30) improve, there is a parallel need for automated, robust analytical approaches. The mathematical field of topological data analysis provides a powerful framework for structural analysis of SMLM data. We have introduced tools using both persistence based clustering (13) and persistent homology (16) to quantify biological nano-structure. ToMATo provides superior segmentation performance over commonly used algorithms for SMLM data when either structures vary in density, or are close together (6, 9–11). We argue that biological nano-structure is rarely simple and for many applications these conditions will be commonplace, as we show for clustering of collagen receptors in platelets. Moreover selection of free parameters is less sensitive for persistence based clustering, and can be guided by ToMATo diagrams. Whilst Bayesian approaches (12) to parameter setting are very powerful when the underlying nano-structure can be modeled in advance, it is not suitable for exploratory applications where the nano-structure is not known *a priori*. The core concepts in this work could also be transferred to other clustering approaches which could be extended to threshold on persistence (14, 15). For example Voronoï tesselation could be used to calculate the density estimate. The dual graph of the tessellation could also be used to link detections resulting in an algorithm with only one free parameter, providing an interesting avenue for further research.

Persistent homology is designed to reveal topological structure within pointillist datasets and has natural applications for the analysis of SMLM data, as also demonstrated in parallel work (31). Our novel framework moves beyond per cluster statistics such as area and density to reveal new information about the underlying topological features within segmented nano-structures. Importantly, this is a multi-scale approach, revealing unbiased structural information at a range of distance scales simultaneously. Validation of these approaches was performed using nuclear pore and endocytic site proteins. Similar structural information can be gained from SMLM data by image based particle averaging (4, 27). However persistent homology can be used in cases where particle averaging methods may fail, for example, where the underlying structures vary in shape or size but maintains a constant topology. Persistent homology can also quantify the structure of individual clusters, not just average ensembles, and is suitable for studying smaller datasets. Finally image-based analysis workflows are typically complex with many pre-processing steps specific to the application. Our streamlined workflow requires minimal adaptation for different datasets as demonstrated throughout this study.

Our methods for persistence based clustering and persistent homology have been implemented and validated in both 2D and 3D. This framework is applicable for a wide range of problems, revealing new topological information and unprecedented insight into biological nano-structure not attainable from existing tools.

## Methods

### ToMATo clustering

Detection density estimates were calculated by counting the number of other detections within a specified radius. This was implemented using the R package dbscan which uses the C++ library ANN to employ a k-d tree framework for efficient computation. Density modes were calculated using a hill climbing approach and a Rips graph (19). Modes were then merged or designated as noise based on a specified persistence threshold. This was implemented by adapting C++ scripts (GPLv3) described in (13). C++ functions were incorporated into RSMLM using the R package Rcpp. For algorithm comparison the search radius ranged from 1 to 50nm, and the persistence threshold from 0 to 50 detections.

### Other clustering algorithms

Density based spatial clustering of applications with noise (DBSCAN) was implemented using the R package dbscan (11). Edge detections were included in clusters. For algorithm comparison the search radius ranged from 1 to 50nm, and the density threshold from 1 to 100nm.

Ripley K based clustering was implemented by finding the number of detections within a specified search radius, *r*, and using this to calculate the Ripley L function, *L*(*r*), for each detection. *L*(*r*) – *r* was subsequently used as a density estimate and thresholded (6). Filtered detections were grouped into clusters by finding the connected components of the graph formed by linking all filtered detections within *r* (12). For algorithm comparison the search radius, *r*, ranged from 1 to 50nm, and the *L*(*r*) –*r* threshold from −10 to 30.

Voronoï tesselations were calculated using the R package deldir. Tesselation based clustering was implemented by removing all tiles where the normalised detection density was below a specified threshold. Density was either defined as the inverse of tile area (Voronoï zero) (10), or the inverse of mean first rank tile area (Voronoï 1st) (9). Mean first rank tile area is defined as the mean area of the specified tile and all adjacent tiles. Density estimates were normalised by dividing by the mean density across the full field of view. After removing tiles from the tessellation, clusters were formed from the connected components of the dual graph (the Delaunay triangulation). For algorithm comparison the normalised density thresholds ranged from 1 to 5.

### Persistent homology

Rips filtrations and persistence diagrams were computed using the R package TDA which employs the C++ library GHUIDI (32, 33). When the density of points in a persistence diagram was too high for a simple plot, features were grouped into a joint histogram with 1nm^2^ bins and displayed using a heat-map. Visualisations of Rips complexes were created using plex-viewer (https://github.com/atausz/plex-viewer) and the Java library javaPlex (34). Filtration movies were created using POV-Ray.

### Cluster measurements

Cluster area and volume were calculated using the convex hull of all detections in a cluster. Cluster density was defined as the number of cluster detections divided by the cluster area (volume in 3D). The number of holes per cluster was defined as the number of second order topological features in the corresponding persistence diagram above a specified persistence threshold. Similarly, the number of voids corresponds to the number of third order features above a specified persistence threshold.

### Radial averaging of clusters

Clusters were filtered for a specified topology, for example clusters with a single hole. For each filtered cluster the intensity weighted center of mass (COM) was found. For each detection in a cluster the radial distance to the corresponding COM was calculated. To produce the average radial distribution these distances were grouped into 10nm bins (ranging from 0-150nm). To normalise the distribution the value for each bin was divided by the corresponding area 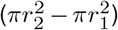, before scaling between 0 and 1. *r*_2_ is the bin maximum, and *r*_1_ the bin minimum.

### dSTORM simulations

Fluorophore blinking characteristics were modeled using a geometric distribution with probability of transition to the dark state set to 0.5 (35). Simulated molecules were bound to an average of 5 fluorophores, randomly distributed between molecules. The localisation uncertainty for each blinking event was determined using a normal distribution centered on the molecule position. Standard deviation for localisation uncertainty was set using a log-normal distribution with mean 2.8 and standard deviation 0.28 (experimentally measured parameters for AlexaFluor647) (36). Detection rate for blinking events was set to 70%. 10% false detections (noise) were added and distributed randomly across the field of view.

### Simulation of data for comparison of clustering algorithms

Simulations for low, high and mixed (a mixture of high and low) density clusters were generated, either in close proximity or well separated. For each of these six scenarios four Gaussian clusters were placed on a field of view of size 400nm^2^ for well separated clusters, 300nm^2^ otherwise. The standard deviation for low and high density clusters was set to 5nm and 20nm respectively. For mixed density simulations two high density and two low density clusters were generated. For well separated simulations all clusters were separated by 200nm. Otherwise low and high density clusters were separated by 100nm and 60nm respectively. After generating molecule locations the dSTORM imaging process was simulated. Simulations were repeated twenty times and algorithm performance, for a given parameter set, was averaged across all repeats.

### Simulation of nano-structures for topological analysis

For ring simulations 40 molecules were even spaced on the circumference of a circle with radius 60nm. For hollow sphere simulations 100 molecules were randomly placed on the surface of a sphere with radius 75nm. The standard deviation for Gaussian clusters was set to the ring (or sphere) radius / 1.5 (87% of molecules will lie within the radius). For all scenarios five clusters were distributed on a 1*μ*m^2^, or 1*μ*m^3^, field of view according to a uniform random distribution, with re-selection if closer to another cluster than the diameter of the ring (or sphere). For randomly distributed simulations molecules were placed on the field of view according to complete spatial randomness. There where 200 and 500 molecules per field of view for the 2D and 3D simulations respectively. After generating molecule locations the dSTORM imaging process was simulated. Simulations were repeated twenty times for each scenario.

### Platelet preparation and spreading

Human washed platelets were prepared from blood samples donated by healthy, consenting volunteers (local ethical review no: ERN-11-0175). Blood was drawn via venipuncture into sodium citrate as the anticoagulant and then acid/citrate/dextrose (ACD) added to 10% (v:v). Blood was centrifuged at 200xg for 20 minutes. Platelet rich plasma (PRP) was removed and then centrifuged at 1000×g for 10 min in the presence of 0.1 *μ*g/ml prostacyclin. Plasma was removed and the platelet pellet was resuspended in modified Tyrode’s buffer (134mM NaCl, 0.34mM Na_2_HPO_4_, 2.9mM KCl, 12mM NaHCO_3_, 20mM HEPES, 5mM glucose, 1mM MgCl_2_; pH 7.3) containing ACD and 0.1*μ*g/ml prostacyclin before being centrifuged for 10 min at 1000×g. The washed platelet pellet was resuspended in modified Tyrode’s buffer, left to rest for 30 min and the platelet count adjusted to 2*×* 107 platelets/ml.

### Platelet spreading and immunolocalisation

Glass-bottom MatTek dishes were coated with 10*μ*g/ml Horm collagen (diluted in manufacturer supplied diluent; Takeda, UK) overnight at 4°C before being blocked in 5mg/ml BSA for 1 hour at room temperature. Washed platelets were added to the dishes and allowed to spread for 45 minutes at 37°C before the addition of the Syk inhibitor PRT060318 (10*μ*M) or DMSO control, each diluted in modified Tyrode’s buffer. The platelets were returned to 37°C for a further 15 minutes before being washed once in PBS and then fixed in 10% formalin solution for 10 min. Following PBS washes the platelets were permeabilised with 0.1% Triton X-100 for 5 min and then washed in PBS and blocked for 30 min in block buffer (1% BSA, 2% goat serum in PBS). The platelet integrin *α*2*β*1 was immuno-labelled with 5*μ*g/ml anti-CD49b (clone 16B4; AbD Serotec) diluted in block buffer for 1 hour at room temperature. Platelet integrin *α*2*β*1 was secondary labelled with anti-mouse-Alexa647 and F-actin was labelled with phalloidin-Alexa488 (both Invitrogen and diluted 1:300 in block buffer). Platelets were washed and stored in PBS.

### dSTORM imaging of platelet integrin *α2β1*

Labelled platelets were imaged on a Nikon N-STORM system using a 100 × 1.49NA TIRF objective. The system contains a Ti-E stand with Perfect Focus, a Andor IXON Ultra 897 EMCCD camera and a Agilent Ultra High Power Dual Output Laser bed (containing a 170mW 647nm laser and a 20mW 405nm laser). DIC and TIRF images of the platelets were taken to identify areas containing platelets and collagen fibres. To induce fluorophore blinking of the Alexa647 labelled integrin, platelets were imaged in a PBS-based buffer containing 100 mM MEA, 50ug/ml glucose oxidase and 1ug/ml catalase, pH 7.5, as detailed in (37). For each image, 20,000 frames were captured using NIS Elements 4.2 with an exposure time of 20ms, gain 300 and conversion gain 3.

Detections were localised using the ThunderSTORM plugin for Fiji with a Gaussian PSF model and maximum likelihood fitting (38). Image visualisations were produced with the normalised Gaussian method.

### Nup107 and endocytic protein datasets

2D and 3D SMLM Nup107 data was recently published and described in (25) where Nup107–SNAP–Alexa Fluor 647 was imaged using dSTORM in U-2 OS cells. 3D localiazation was achieved using an experimental PSF model.

SMLM datasets for Las17, Ede1 and Sla1 were recently described in (4). In short endocytic proteins were endogenously tagged with the photoconvertible protein mMaple (39) in budding yeast strains and imaged in high-throughput using a homebuilt SMLM imaging system (40).

### Detection filtering

Detections in the integrin *α*2*β*1 and Nup107 datasets were filtered by intensity with a minimum value of 1000 photons. For all real data detections that were found in consecutive frames within a distance of 75nm were grouped into a single detection.

### Code availability

RSMLM has been released under the GNU General Public License v3.0 and is available at https://github.com/JeremyPike/RSMLM. Tutorials for this library implemented as Binder ready Jupyter notebooks are available at https://github.com/JeremyPike/RSMLM-tutorials.

## ACKNOWLEDGEMENTS

The authors gratefully acknowledge financial support from the Centre of Membrane Proteins and Receptors (COMPARE), Universities of Birmingham and Nottingham, Midlands, UK; and from the British Heart Foundation through the Chair award (CH0/03/003) to Steve P Watson, who also provided many useful comments.

## Supplementary Note 1: Supplementary Figures and Tables

**Fig. S1.**
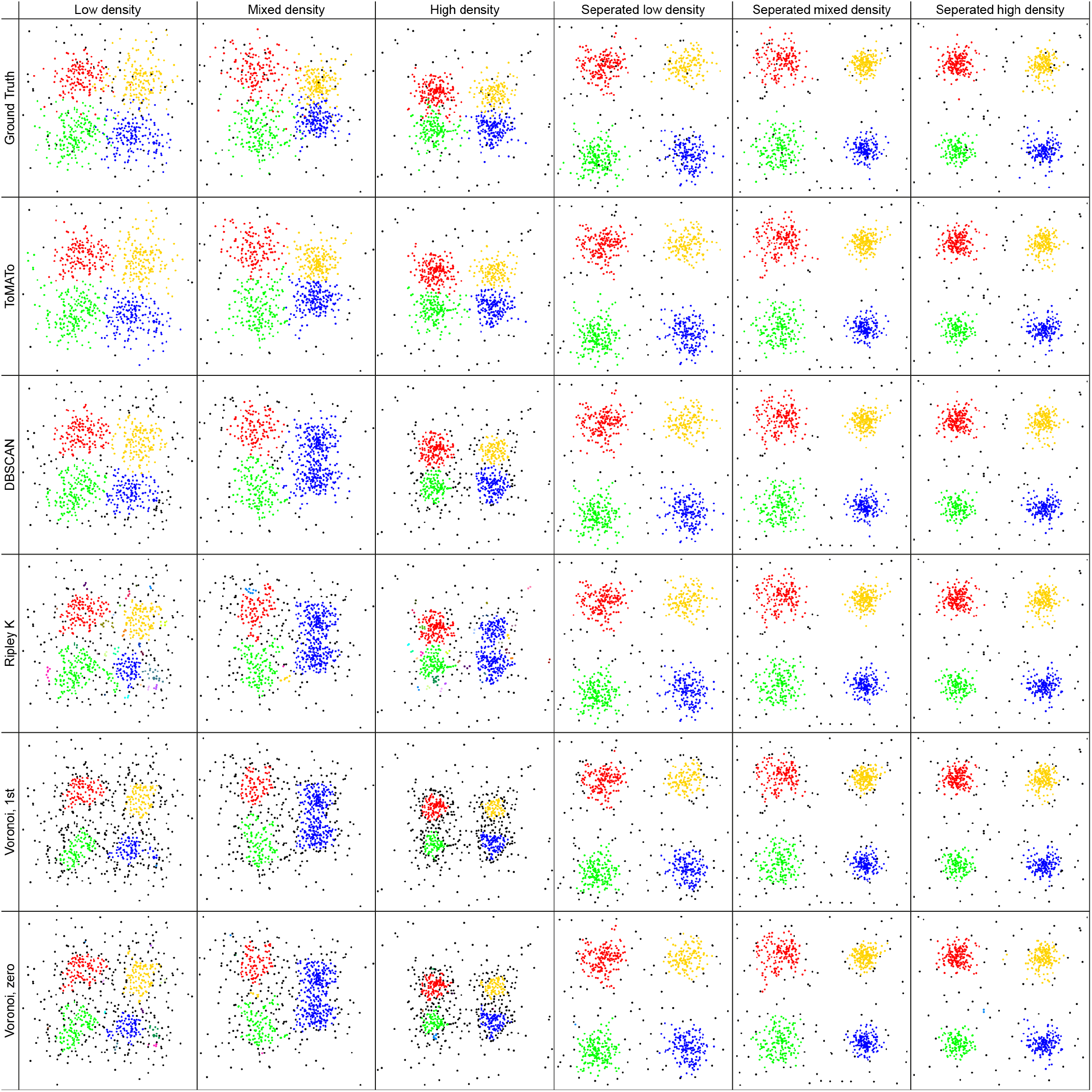
Example cluster results. For each algorithm and scenario the parameters which produced the highest mean performance were used.

**Fig. S2.**
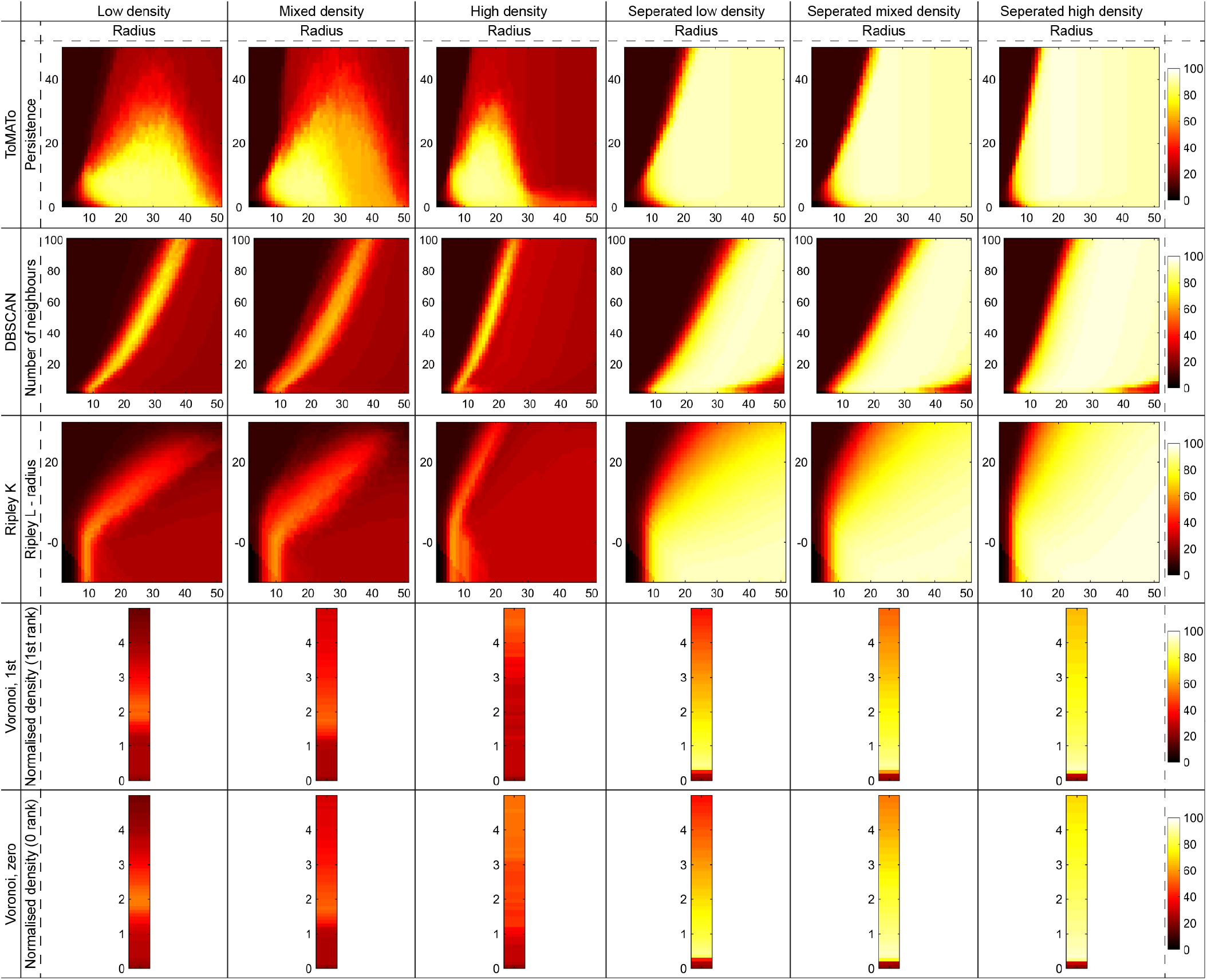
Heatmaps showing the performance of each clustering algorithm across all clustering scenarios and parameter sets. Performance is defined as the percentage of correctly assigned detections and was averaged across twenty simulations.

**Fig. S3.**
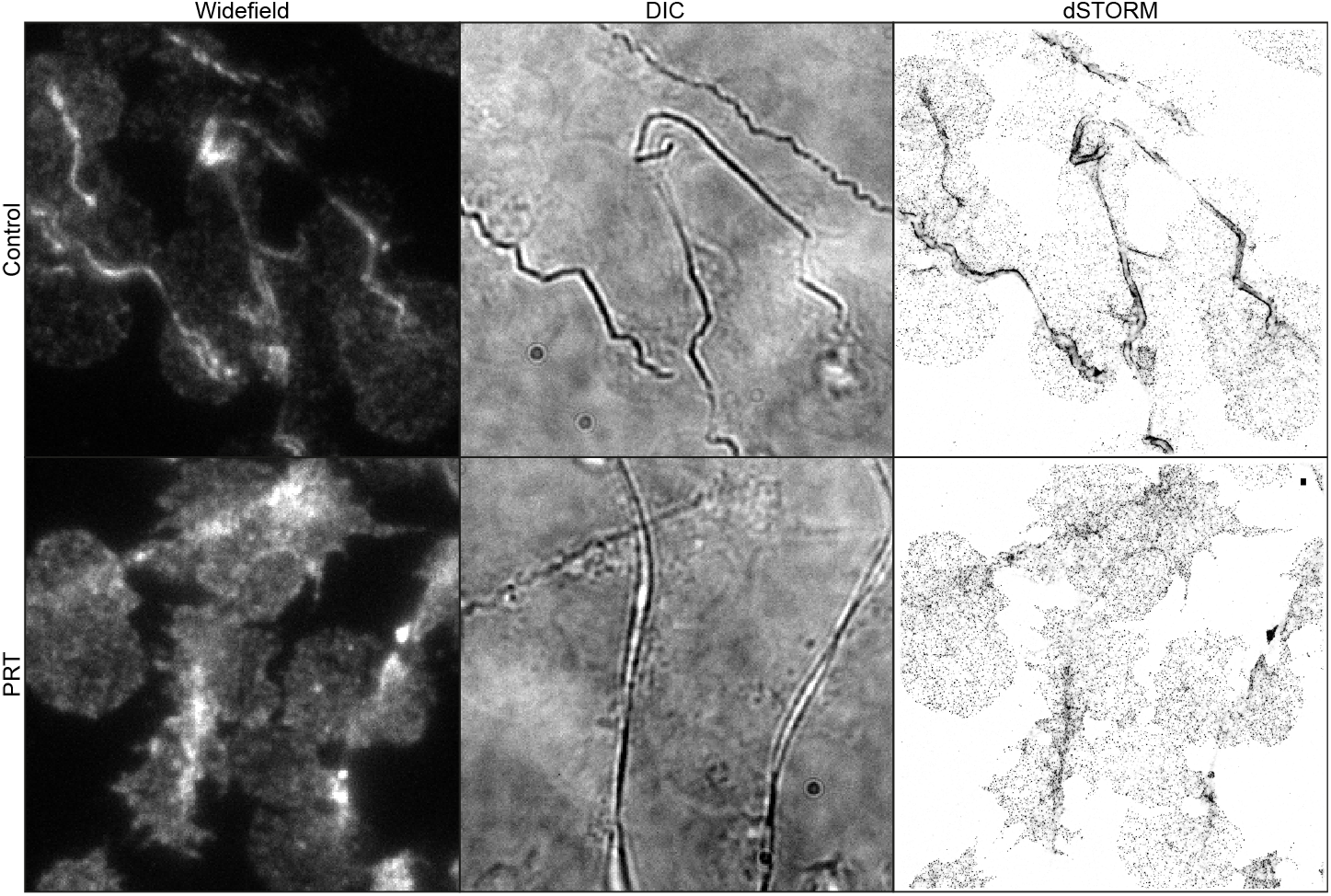
Imaging of integrin *α*2*β*1 by dSTORM. Platelets were seeded on collagen fibers and treated either with PRT060318, or DMSO (control). Representative widefield, DIC and dSTORM images are shown. Collagen fibers can be seen in the DIC image. Scale-bar 500nm.

**Fig. S4.**
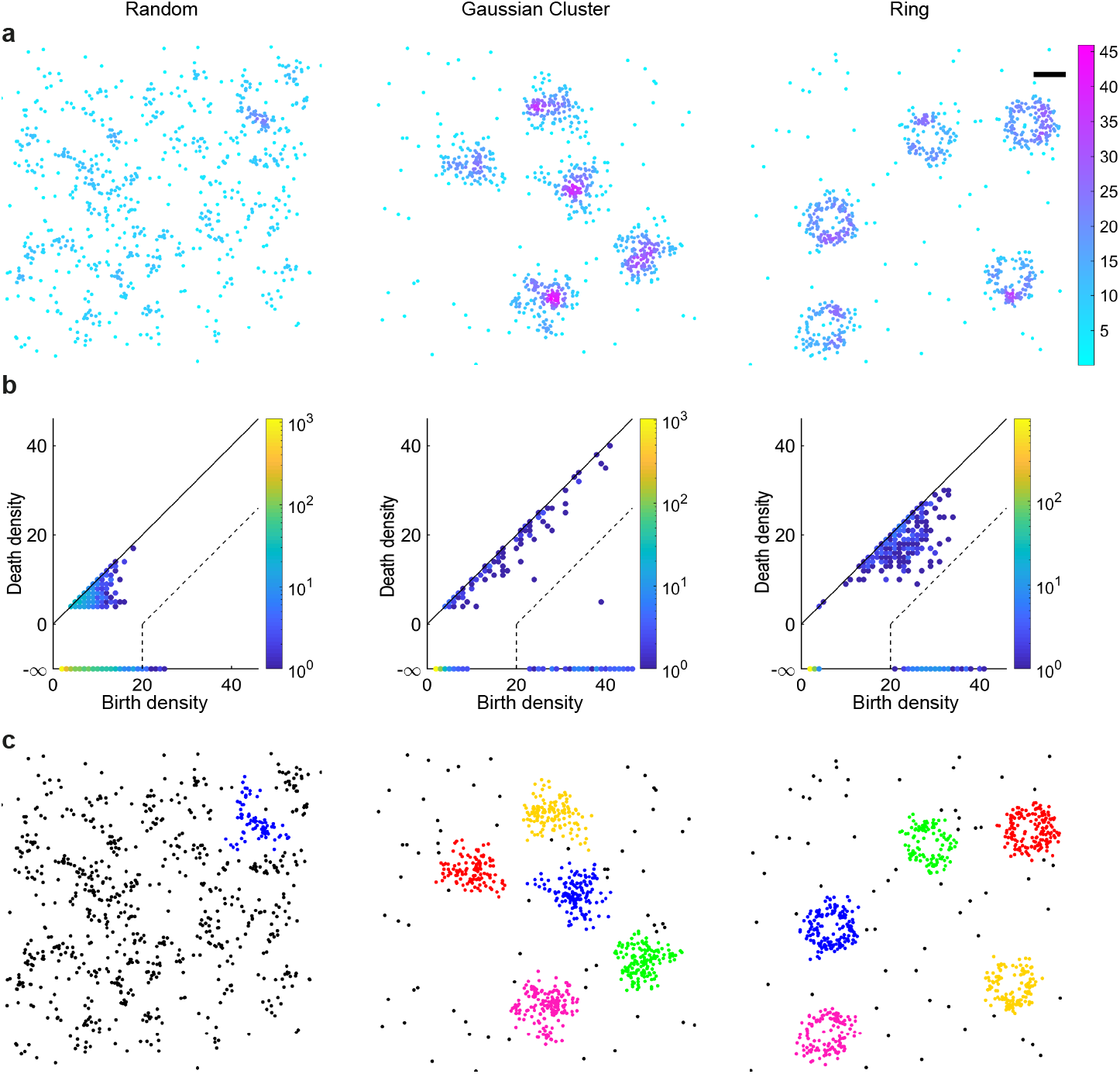
Persistence based clustering as a pre-processing step for topological analysis. (**a**) Detection density for simulations of randomly distributed molecules, Gaussian clusters and rings were estimated by counting the number of other detections within 30nm. Scale-bar 100nm. (**b**) ToMATo diagrams showing the birth and death coordinates for all candidate clusters. For unbiased visualisation clusters across repeated simulations have been grouped. A persistence threshold of 20nm was used to merge clusters (dotted line). (**c**) Final clustering results after merging. Noise points are shown in black.

**Fig. S5.**
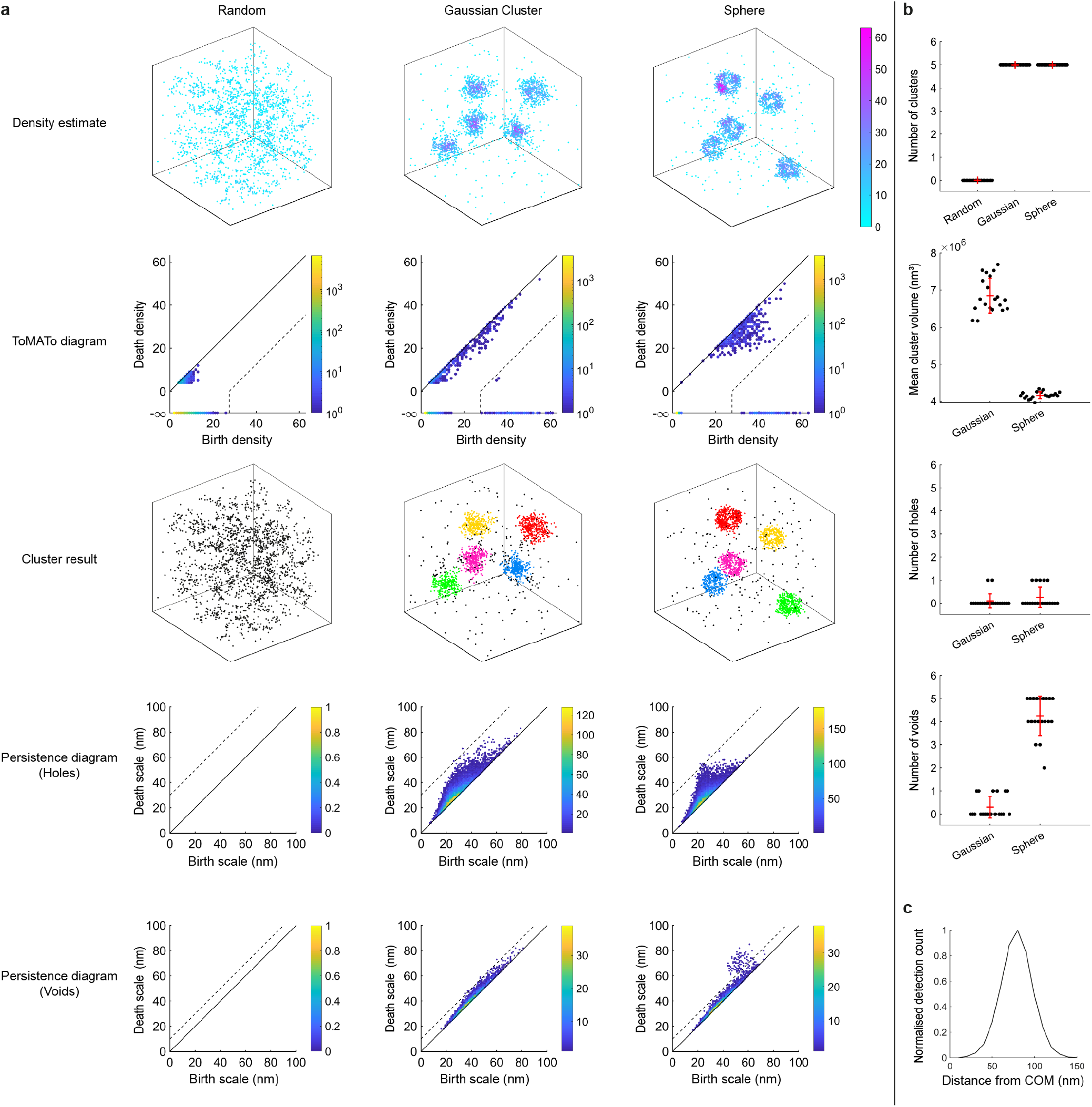
Persistence based clustering and persistent homology for analysis of 3D SMLM datasets. (**a**) Example simulations for randomly distributed molecules, Gaussian clusters and hollow spheres with radius 75nm. Detection density was estimated by counting the number of other detections within 40nm. ToMATo diagrams were used to select a persistence threshold for merging of density clusters (27.5 detections, dotted line). After cluster merging persistent homology was performed to produce persistence diagrams for both 2D (holes) and 3D (enclosed voids) features. Features from all clusters were grouped into a single diagram per scenario and dimension. Persistence thresholds of 30nm and 10nm were selected for holes and voids respectively. (**b**) Number of clusters, mean cluster area, number of holes and number of voids. The number of voids is higher for the hollow sphere simulation and there was no significant difference in the number of holes between the Gaussian and sphere simulations. This is consistent with the topology of the simulated structures as a hollow sphere has a single enclosed void but no holes. As expected the cluster area is lower for the sphere simulation. Error bars are mean ± s.d.. (**c**) Averaged radial distribution for all clusters with a single void from the spherical simulation. Peak at 80nm.

**Fig. S6.**
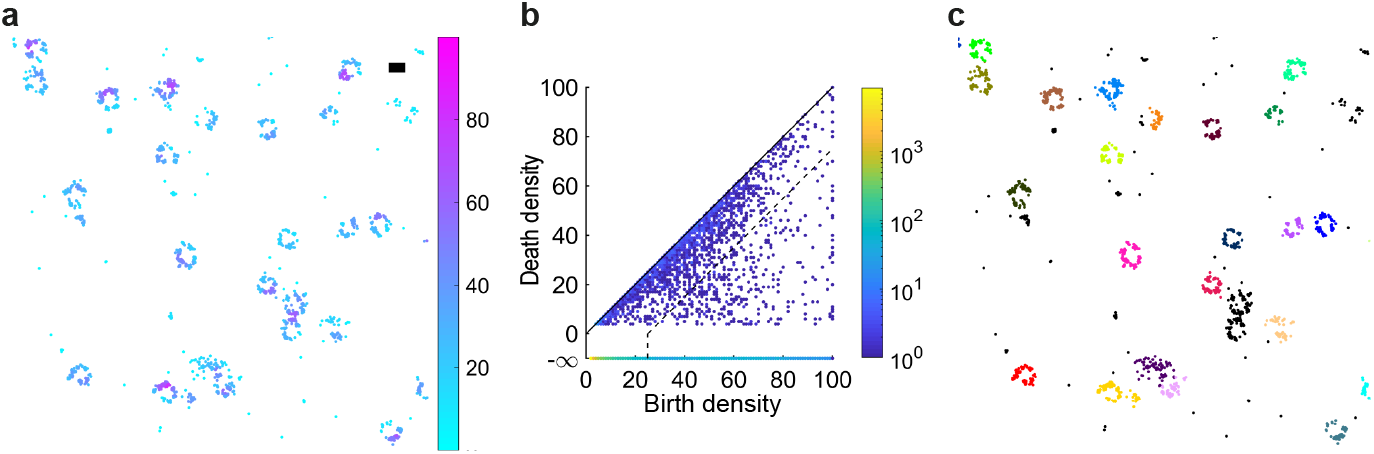
Persistence based clustering of nuclear pore component Nup107 in 2D. (**a**) Density estimate was calculated using a search radius of 25nm. Representative cropped field of view shown. Scale-bar 100nm. (**b**) ToMATo diagram showing all candidate clusters. Persistence threshold set to 25 detections (dotted line). (**c**) Result of ToMATo clustering to segment nuclear pore complexes. Clusters were filtered by number of detections (20 *−* 400) and area (*π ×* 40^2^ *−π ×* 100^2^nm^2^).

**Fig. S7.**
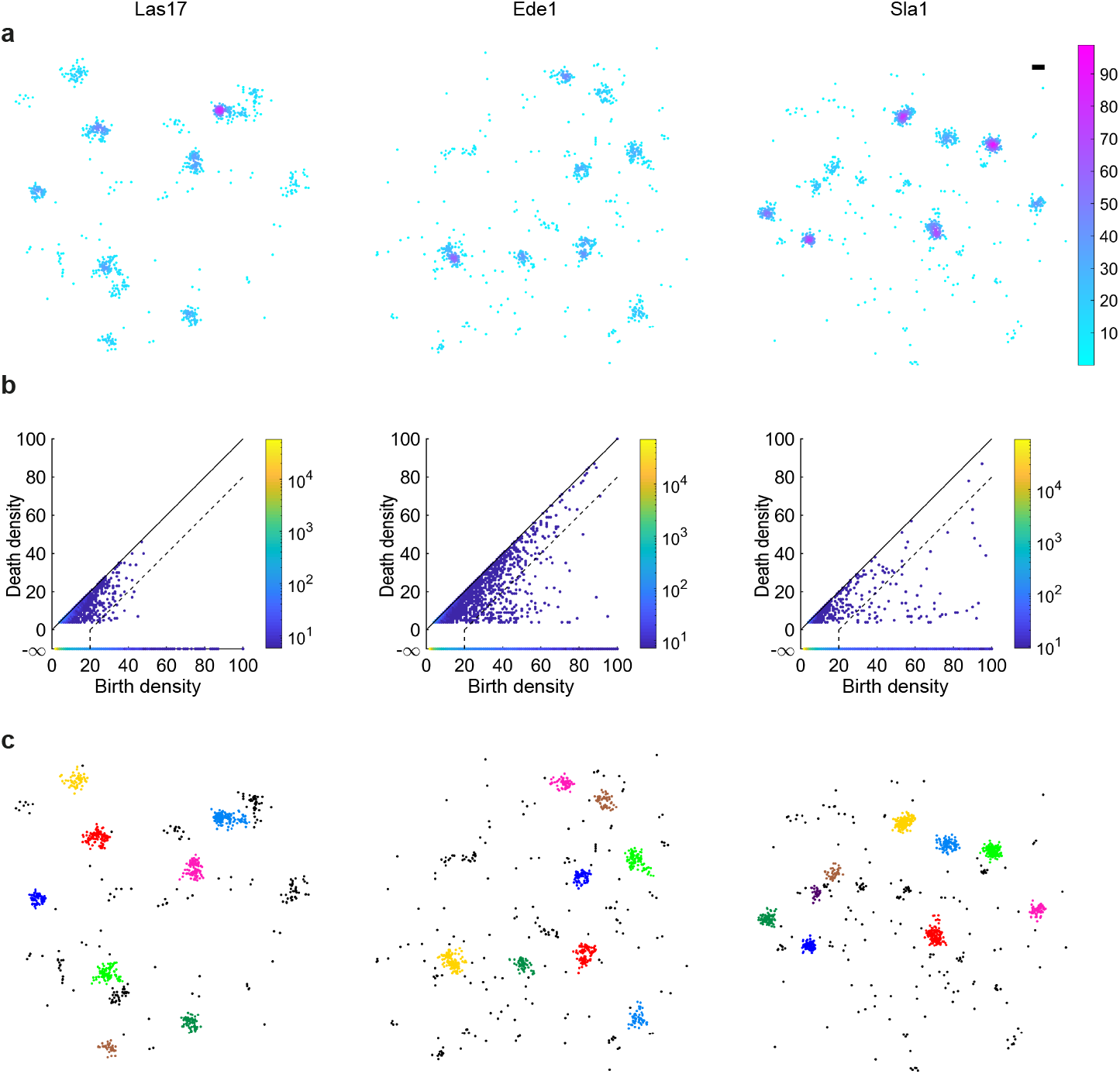
Persistence based clustering of endocytic proteins; Las17, Ede1 and Sla1. (**a**) Density estimates were calculated using a search radius of 40nm. Representative cropped field of view shown. Scale-bar 100nm. (**b**) ToMATo diagrams showing candidate clusters. Persistence threshold set to 20 detections (dotted lines). All fields of view grouped into a single diagram per condition. (**c**) Result of ToMATo clustering to segment endocytic sites. Clusters were filtered by number of detections (20+) and area (*π* × 30^2^ − *π* × 130^2^nm^2^).

**Fig. S8.**
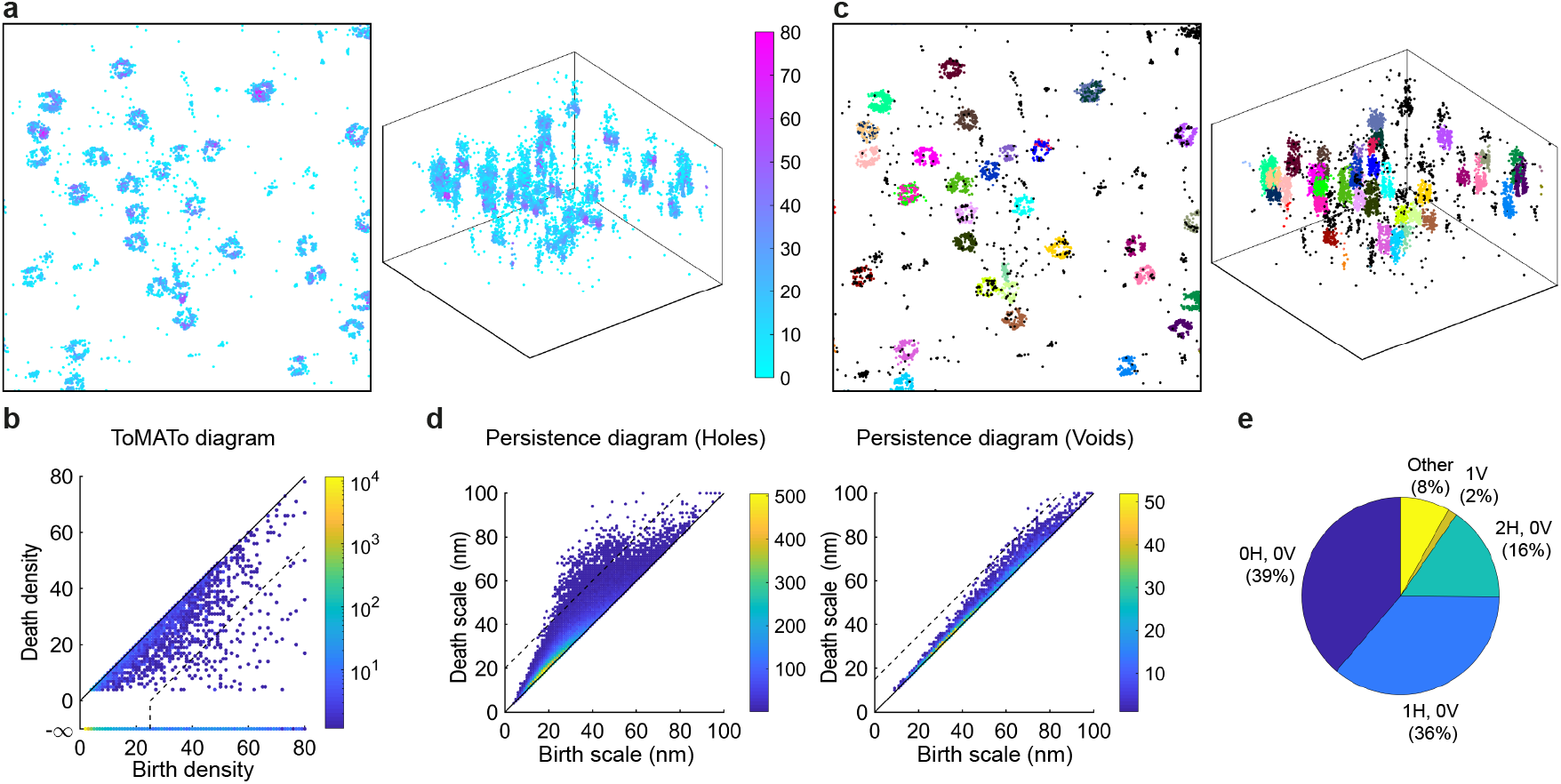
Clustering and topological analysis of nuclear pore component Nup107 in 3D. (**a**) Density estimate was calculated using a search radius of 50nm. Representative cropped field of view shown as a projection and 3D scatter-plot. (**b**) ToMATo diagram where a persistence threshold of 25nm was applied (dotted line). (**c**) Result of ToMATo clustering to segment nuclear pore complexes in 3D. Clusters were filtered by number of detections (20 – 400) and volume (4*/*3*π* × 40^3^ – 4*/*3*π* × 150^3^nm^3^). (**d**) Persistence diagrams for 2D and 3D topological features. Thresholds of 20nm and 15nm were chosen for holes and voids respectively. (**e**) Percentage of clusters with a given topological configuration. Specified in terms of number of holes (H) and voids (V) per cluster.

